# Nanopore sequencing of long ribosomal DNA amplicons enables portable and simple biodiversity assessments with high phylogenetic resolution across broad taxonomic scale

**DOI:** 10.1101/358572

**Authors:** Henrik Krehenwinkel, Aaron Pomerantz, James B. Henderson, Susan R. Kennedy, Jun Ying Lim, Varun Swamy, Juan Diego Shoobridge, Nipam H. Patel, Rosemary G. Gillespie, Stefan Prost

**Author notes:** Corresponding authors: Henrik Krehenwinkel and Stefan Prost.

## Abstract

**Background:** In light of the current biodiversity crisis, DNA barcoding is developing into an essential tool to quantify state shifts in global ecosystems. Current barcoding protocols often rely on short amplicon sequences, which yield accurate identification of biological entities in a community, but provide limited phylogenetic resolution across broad taxonomic scales. However, the phylogenetic structure of communities is an essential component of biodiversity. Consequently, a barcoding approach is required that unites robust taxonomic assignment power and high phylogenetic utility. A possible solution is offered by sequencing long ribosomal DNA (rDNA) amplicons on the MinION platform (Oxford Nanopore Technologies).

**Results:** Using a dataset of various animal and plant species, with a focus on arthropods, we assemble a pipeline for long rDNA barcode analysis and introduce a new software (MiniBar) to demultiplex dual indexed nanopore reads. We find excellent phylogenetic and taxonomic resolution offered by long rDNA sequences across broad taxonomic scales. We highlight the simplicity of our approach by field barcoding with a miniaturized, mobile laboratory in a remote rainforest. We also test the utility of long rDNA amplicons for analysis of community diversity through metabarcoding and find that they recover highly skewed diversity estimates.

**Conclusions:** Sequencing dual indexed, long rDNA amplicons on the MinION platform is a straightforward, cost effective, portable and universal approach for eukaryote DNA barcoding. Long rDNA amplicons scale up DNA barcoding by enabling the accurate recovery of taxonomic and phylogenetic diversity. However, bulk community analyses using long-read approaches may introduce biases and will require further exploration.

## Introduction

The world is changing at an unprecedented rate, threatening the integrity of biological communities [1, 2]. To understand the impacts of change, whether a system is close to a regime shift, and how to mitigate the impacts of a given environmental stressor, it is important to consider the biological community as a whole. In recognition of this need, there has been a shift in emphasis from studies that focus on single indicator taxa, to comparative studies across multiple taxa and metrics that consider the properties of entire communities [3]. Such efforts require accurate information on the identity of the different biological entities within a community, as well as the phylogenetic diversity that they represent.

Comparative ecological studies across multiple taxa have been greatly simplified by molecular barcoding [4], where species identifications are based on short PCR amplicon “barcode” sequences. Different barcode marker genes have been established across the tree of life [5, 6], with mitochondrial cytochrome oxidase subunit I (COI) commonly used for animal barcoding [4]. The availability of large sequence reference databases and universal primers, together with its uniparental inheritance and fast evolutionary rate, make COI a useful marker to distinguish even recently diverged taxa. In recent years, DNA barcoding has greatly profited from the emergence of next generation sequencing (NGS) technology. Current NGS platforms enable the parallel generation of barcodes for hundreds of specimens at a fraction of the cost of Sanger sequencing [7]. Furthermore, NGS technology has enabled metabarcoding, the sequencing of bulk community samples, which allows scoring the diversity of entire ecosystems [8].

However, despite their undeniable advantages, barcoding approaches using short, mitochondrial markers have several drawbacks. The phylogenetic resolution offered by short barcodes is very limited, as they contain only a restricted number of informative sites. This problem is exacerbated by the fast evolutionary rate of mitochondrial DNA, which leads to a quick saturation with mutations, increasing the probability of homoplasy between divergent lineages. The accurate estimation of phylogenetic diversity across wide taxonomic scales, however, is an important component of biodiversity research [9]. Moreover, mitochondrial DNA is not always the best marker to reflect species differentiation, as different factors are known to inflate mitochondrial differentiation in relation to the nuclear genomic background. For example, male biased gene flow [10] or infections with reproductive parasites [11] (e.g. *Wolbachia)* can lead to highly divergent mitochondrial lineages in the absence of nuclear differentiation. In contrast, introgressive hybridization can cause the complete replacement of mitochondrial genomes (see e.g. [12, 13]), resulting in shared mitochondrial variation between species.

Considering this background, it would be desirable to complement mitochondrial DNA based barcoding with additional information from the nuclear genome. An ideal nuclear barcoding marker should possess sufficient variation to distinguish young species pairs, but also provide support for phylogenetic hypotheses between divergent lineages. Moreover, the marker should be present across a wide range of taxa and amplification should be possible using universal primers. A marker that fulfils all the above requirements is the nuclear ribosomal DNA (rDNA).

As an essential component of the ribosomal machinery, rDNA is a common feature across the tree of life from microbes to higher eukaryotes [14]. All eukaryotes share homologous transcription units of the 18S, 5,8S and 28S-rDNA genes, which include two internal transcribed spacers (ITS1 and ITS2) [15]. Due to varying evolutionary constraints acting on different parts of the rDNA, it consists of regions of extreme evolutionary conservation, which are interrupted by highly variable sequence stretches [16]. While some rDNA gene regions are entirely conserved across all eukaryotes, the two ITS sequences are distinguished by such rapid evolutionary change that they separate even lineages within species [5, 17]. rDNA markers thus offer taxonomic and phylogenetic resolution at a very broad taxonomic scale. As an essential component of the translation machinery, nuclear rDNA is required in large quantities in each cell. It is thus present in multiple copies across the genome [15] and is readily accessible for PCR amplification. Due to the above advantages, rDNA already is a popular and widely used marker for molecular taxonomy and phylogenetics in many groups of organisms [5, 6, 15, 17, 18].

Spanning about 8 kb, the ribosomal cluster is fairly large, and current barcoding protocols, e.g. using Sanger sequencing or Illumina amplicon sequencing, can only target short sequence stretches of 150 - 1,000 bp. Such short stretches of 28S and 18S are often too conserved to identify young species pairs [19]. The ITS regions, on the other hand, are too variable to design truly universal primers, leading to a considerable amount of taxon dropout during PCR. Moreover, ITS sequences can show considerable length variation between taxa, and are often too long for short amplicon-based barcoding [20]. Consequently, it would be ideal to amplify and sequence a large part of the ribosomal cluster in one fragment. A solution to sequence the resulting long amplicons is offered by recent developments in third generation sequencing platforms, which now enable researchers to generate ultra-long reads, of up to 800 kb [21]. Recently, amplicons of several kilobases of the rDNA cluster were sequenced using Pacific Bioscience (PacBio) technology, to explore fungal community composition [22, 23]. But while PacBio sequencing is well suited for long amplicon sequencing, it is currently not readily available to every laboratory due to the high cost and limited distribution of sequencing machines.

A cost-efficient and readily available alternative is provided by nanopore sequencing technology. The MinION sequencer (Oxford Nanopore Technologies) is small in size, lightweight, allows for sequencing of several Gb’s of DNA with average read lengths over 10 kb on a single flow cell [24] and is available starting at $1,000. Despite a raw read error rate of about 12-22 % [21], highly accurate consensus sequences can be called from nanopore data [25, 26]. The MinION is well suited for amplicon sequencing, and a simple dual indexing strategy can be used to demultiplex amplicon samples [27]. This technology offers tremendous potential for long-read barcoding applications, as recently shown in an analysis in fungi [26]. However, current analyses are still exploratory or limited in taxonomic focus and streamlined analysis pipelines to establish the method across the eukaryote tree of life are still missing.

Considering this background, we explore the feasibility of nanopore sequencing of long rDNA amplicons as a simple, cost efficient and universal eukaryote DNA barcoding approach. We compile a workflow from PCR amplification, to library preparation, to demultiplexing and consensus calling (see Fig. 1 for an overview). We explore the error profile of nanopore consensus sequences and introduce MiniBar, a new software to demultiplex dual indexed nanopore amplicon sequences. We test the utility of the ribosomal cluster for molecular taxonomy and phylogenetics across divergent plant and animal taxa. A particular focus of our analysis are arthropods, the most diverse group in the animal kingdom [28], which are highly threatened by current mass extinctions [29]. Using a dataset of spiders, we compare the taxonomic resolution of the ribosomal cluster with that offered by molecular barcoding using mitochondrial COI, the currently preferred barcode marker for arthropods. Oxford Nanopore Technologies’ MinION is a portable sequencer, and Nanopore based DNA barcoding has been applied in remote sites outside of conventional labs (see eg. [25, 30, 31]). Such field-based applications confront researchers with additional complexities and challenges. To highlight the simplicity of our approach, we tested it under field conditions and generated long rDNA barcode sequences using a miniaturized mobile laboratory in a Peruvian rainforest.

**Figure 1.**
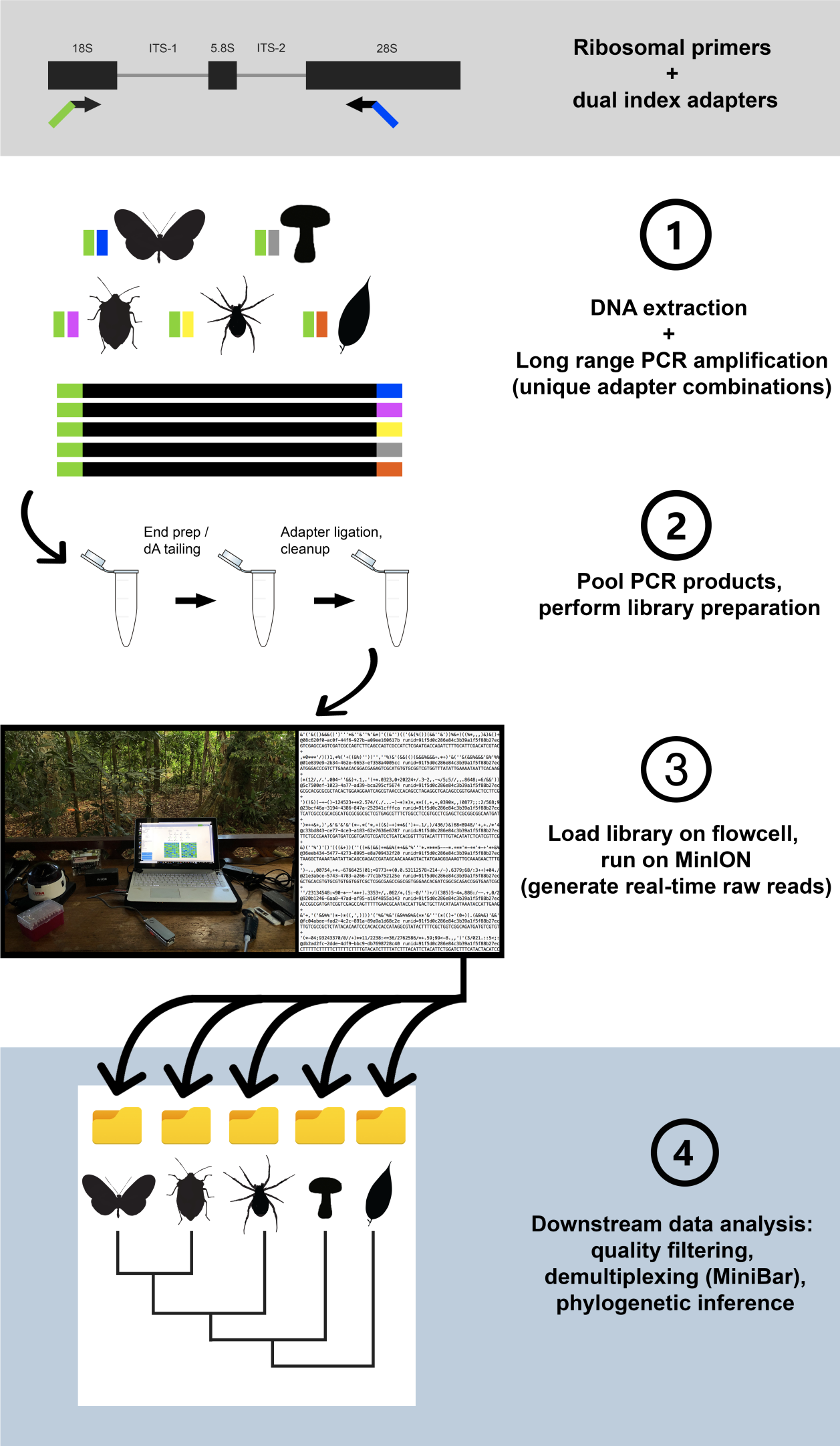
Workflow for the design, amplification, and sequencing of the ribosomal DNA cluster.

We also tested the efficacy of long-read rDNA sequencing for metabarcoding of bulk community samples. A study of bacterial communities [32] suggests Nanopore long-read sequencing as a powerful tool for community characterization, but also found pronounced biases in the recovered taxon abundance. Currently, little is known about the utility of long-read sequencing for animal community analysis. Metabarcoding protocols for community samples need to be carefully optimized, as they can suffer from pronounced taxonomic biases, e.g. due to primer binding or polymerase efficiency [33]. Well established Illumina based short read metabarcoding protocols can account for these biases and allow for a very high qualitative and even quantitative recovery of taxa in communities [34]. However, additional, yet unexplored, biases may affect long-read metabarcoding. We thus also test the utility of long-read rDNA barcoding to recover taxonomic diversity from arthropod mock communities. We compare the qualitative (species richness) and quantitative (species abundance) recovery of taxa by long-read sequencing with that based on short read Illumina amplicon sequencing of the 18SrDNA.

Overall, we demonstrate that long rDNA amplification and sequencing on the MinION platform is a straightforward, cost effective, and universal approach for eukaryote DNA barcoding. It combines robust taxonomic assignment power with high phylogenetic resolution and will enable future analyses of taxonomic and phylogenetic diversity across wide taxonomic scales.

## Materials and Methods

### DNA extraction, PCR and library preparation

We analyzed 114 specimens of eukaryotes including 17 insect and 42 spider species, two annelid and nine plant species (Supplementary Table 1). Some feeder insects and the annelids were purchased at a pet store. The remaining specimens were collected in oak forest on the University of California Berkeley’s campus or in native rainforests of the Hawaiian Archipelago (under the Hawaii DLNR permit: FHM14-349). We particularly focused our arthropod sampling on spiders, which are ubiquitous and essential predators in all terrestrial ecosystems. Recent phylogenomic work [35] provided us with a solid baseline to test the efficiency of rDNA amplicons for phylogenetic and taxonomic purposes. We included a taxonomically diverse collection of 16 spider families from the Araneoidea, the RTA clade and a haplogyne outgroup species. Within spiders, we additionally focused on the genus *Tetragnatha*, which has undergone a striking adaptive radiation on Hawaii.

DNA was extracted from each sample using the Qiagen Archivepure kit (Qiagen, Valencia, CA, USA) according to the manufacturer’s protocols. The DNA integrity was checked on an agarose gel. Only samples with high DNA integrity were used for the following PCRs. All DNA extracts were quantified using a Qubit fluorometer using the high sensitivity dsDNA assay (Thermo Fisher, Waltham, MA, USA) and diluted to concentrations of 20 ng/μl. We designed a primer pair of each 27 bases to amplify a ~4,000 bp fragment of the ribosomal DNA, including partial 18S and 28S as well as full ITS1, 5.8S and ITS2 sequences (18S_F4 GGCTACCACATCYAARGAAGGCAGCAG and 28S R8 TCGGCAGGTGAGTYGTTRCACAYTCCT). The primers were designed using alignments of partial 18S and 28S sequences of ~1,000 species of eukaryotes, with a focus on animals. The primers targeted highly conserved regions across all analyzed taxa. Degenerate sites were incorporated to account for variation. We aimed for high annealing temperatures (65-70°C) to impose stringent amplification. These were calculated using the NEB Tm Calculator (https://tmcalculator.neb.com/#!/main).

To index every PCR amplicon separately, we used a dual indexing strategy with each primer carrying a unique 15 bp index sequence at its 5’-tail. Index sequences were designed using Barcode Generator (http://comailab.genomecenter.ucdavis.edu/index.php/Barcode_generator) with a minimum distance of 10 bases between each index. A total of 15 forward and 16 reverse indexes were designed. Every sample was amplified separately using the Q5 Hot Start High-Fidelity 2X Master Mix (NeB, Ipswitch, MA, USA) in 15 μ! reactions, at 68°C annealing temperature, with 35 PCR cycles and using 50 ng of template DNA per PCR. All PCR products were checked and quantified on an agarose gel and then pooled. The final pool was cleaned from residual primers by 0.75 X AMpure Beads XP (Beckman Coulter, Brea, CA, USA). DNA library preparation was carried out according to the 1D PCR barcoding amplicons SQK-LSK108 protocol (Oxford Nanopore Technologies, Oxford, UK). Barcoded DNA products were pooled with 5 μl of DNA CS (a positive control provided by ONT) and an end-repair was performed (NEB-Next Ultra II End-prep reaction buffer and enzyme mix), then purified using AMPure XP beads. Adapter ligation and tethering was carried out with 20 μl Adapter Mix and 50 μl of NEB Blunt/TA ligation Master Mix. The adapter-ligated DNA library was then purified with AMPure beads XP, followed by the addition of Adapter Bead binding buffer, and finally eluted in 15 μl of Elution Buffer. Each R9 flow cell was primed with 1000 μl of a mixture of Fuel Mix and nuclease-free water. Twelve μl of the amplicon library were diluted in 75 μL of running buffer with 35 μL RBF, 25.5 uL LLB, and 2.5 μL nuclease-free water and then added to the flow cell via the SpotON sample port. The “NC_48Hr_sequencing_FLO-MIN107_SQK-LSK108_plus_Basecaller.py” protocol was initiated using the MinION control software, MinKNOW.

### Field trial in the Amazon rainforest

A field trial using the protocol described above was conducted in Tambopata, Peru, at the Refugio Amazonas lodge (−12.874797, −69.409669) using two butterflies, a grasshopper, one mosquito, unidentified insect eggs and two plant specimens. Collection permits in Peru were issued by the Servicio Nacional Forestal y de Fauna Silvestre, 403-2016-SERFOR-DGGSPFFS, 019-2017-SERFOR-DGGSPFFS. DNA extractions, PCR and library preparation were performed in the field using a highly miniaturized laboratory consisting of portable equipment. Equipment used for sequencing under remote tropical conditions is described in further detail in Pomerantz, et al. [25]. DNA extractions were carried out with the Quick-DNA Miniprep Plus Kit (Zymo Research, Irvine, CA, USA) according to manufacturer’s protocol. PCRs were performed using the Q5 Hot Start High-Fidelity 2X Master Mix and the same primers as described above. A battery operated portable miniPCR device (Amplyus, Cambridge, MA, USA) was used to run PCRs. The sequencing on the MinION was carried out as described above.

## Bioinformatics

### Raw data processing and consensus calling

The fastq files generated by the ONT software MinKNOW were de-multiplexed using MiniBar (see description below), with index edit distances of 2, 3, and 4 and a primer edit distance of 11. Next, the reads were filtered for quality (>13) and size (>3kb) using Nanofilt [36](https://github.com/wdecoster/nanofilt). Individual consensus sequences were created using Allele Wrangler (https://github.com/transplantation-immunology/allele-wrangler/) for demultiplexed fastq files with a minimum coverage of 30. Error correction was performed using RACON [37] (https://github.com/isovic/racon). To do so, we first mapped all the reads back to the consensus using minimap (https://github.com/lh3/minimap2). We performed two cycles of running minimap and RACON. Final consensus sequences were compared against the NCBI database using BLASTn to check if the taxonomic assignment was correct.

We performed multiple tests to validate and optimize the consensus accuracy of long-read barcode sequences. To comparatively assess the accuracy, we used consensus sequences of short 18S and 28SrDNA amplicons, which were previously generated using Illumina amplicon sequencing for the 47 analyzed Hawaiian *Tetragnatha* specimens (Kennedy unpublished data). These sequences were aligned with the respective stretches of our nanopore consensus sequences using ClustalW in MEGA [38]. All alignments were then visually inspected and edited manually, where necessary. Pairwise distances between Illumina and nanopore consensus were calculated in MEGA.

To measure consensus accuracy over the whole ribosomal amplicon, we utilized genome skimming data [39] for six Hawaiian *Peperomia* plant species (Lim et al unpublished data). 150 bp paired-end TruSeq gDNA shotgun libraries for the six *Peperomia* samples were sequenced on a single HiSeq v4000 lane (Illumina, San Diego, CA, USA). The resulting paired-end reads were trimmed and filtered using Trimmomatic v0.36 [40] and mapped to their respective nanopore consensus sequences using bowtie2 [41] under default parameter values and allowing for minimum and maximum fragment size of 200 and 700 bases respectively. Mapping coverage of Illumina reads to nanopore consensus sequences ranged between 150 - 600 X with a mean of ~ 300 X across all six samples. We called Illumina read based consensus sequences for each *Peperomia* species using bcftools [42], and aligned them with the previously generated nanopore consensus sequences. Pairwise genetic distances were then calculated in MEGA as described above. We performed two independent distance calculations: 1) excluding indels, i.e. only using nucleotide substitutions to estimate genetic distances, and 2) including indels as additional characters.

Our demultiplexing software allows flexible edit distances to identify forward and reverse indexes from Nanopore reads. Due to the high raw read error rate, too large edit distances could lead to carryover between samples during demultiplexing. This carryover could possibly affect the accuracy of the called consensus sequence. On the other hand, too stringent edit distances may result in very large read dropout. Assuming an average error rate of 12-22 %, 3 bp of our 15 bp indexes should maximize sequence recovery. We thus tested index edit distances of 2, 3, and 4 bp in MiniBar for the six *Peperomia* specimens for which we had generated Illumina based consensus sequences. We counted the number of recovered reads and estimated the accuracy of the resulting consensus sequence based on the according edit distances as described above.

A recent study [25] showed that accurate consensus sequences from nanopore data can be generated using only 30x coverage. We tested 18 different assembly coverages from 10 to 800 sequences for a *Peperomia* species, to explore optimal assembly coverage. We randomly subsampled the quality filtered and demultiplexed fastq file for the according specimen each 10 times for each tested assembly coverage. Consensus sequences were then assembled and genetic distances to the Illumina consensus calculated as described above.

### Phylogenetic and taxonomic analysis

We carried out phylogenetic analyses on two hierarchical levels. First, we built a phylogeny for all higher eukaryote taxa in our dataset, which included plants, animals and fungi. Second, we took a closer look into the phylogeny of spiders. The resulting quality checked consensus sequences of all taxa were aligned using ClustalW in MEGA. The alignments were visually inspected and manually edited. Due to the deep divergence in the eukaryote data set, the highly variable ITS sequences could not be aligned and were excluded. For the analyses of spiders, we retained both ITS sequences and aligned the whole rDNA amplicon. Appropriate models of sequence evolution for each gene fragment of the rDNA cluster were identified using PartitionFinder [43]. Phylogenies were built using MrBayes [44], with 4 heated chains, a chain length of 1,100,000, subsampling every 200 generations and a burnin length of 100,000.

Focusing on the endemic Hawaiian *Tetragnatha* species, we also tested the utility of the ribosomal cluster for taxonomic identification, as we also had COI barcodes available for these species. Our dataset contained ribosomal DNA sequences for 47 specimens in 16 species. We calculated pairwise genetic distances between and within all species for the whole ribosomal cluster and for each separate gene region of the rDNA cluster using MEGA. As the 18S and 5.8S did not yield any species level resolution within Hawaiian *Tetragnatha*, they were not analyzed separately. To compare the taxonomic resolution of the ribosomal cluster with that of the commonly used mitochondrial COI, we calculated inter- and intraspecific distances for an alignment of 418 bp of the COI barcode region for the same spider specimens (Kennedy et al. unpublished data). We performed a Mantel test using the R package ade4 [45] to test for a significant correlation between COI and ribosomal DNA based distances. A comparison of intraspecific and interspecific distances for mitochondrial COI and ribosomal DNA also allowed us to test for the presence of a barcode gap.

### Nanopore based arthropod metabarcoding

To test for the possibility of estimating arthropod community composition from Nanopore sequencing, we prepared four mock communities of different amounts of DNA extracts from 9 species of arthropods from different orders (see Supplementary Table 2). The samples were amplified using the Q5 High Fidelity Mastermix as described above at 68 °C annealing temperature and 35 PCR cycles. We additionally tested two variations of PCR conditions. We either reduced the annealing temperature to 63 °C or reduced the PCR cycle number to 25.

In order to compare our results with those from an optimized Illumina short read protocol, we amplified all samples for a ~300 bp fragment of the 18SrDNA using the primer pair 18S2F/18S4R [46]. Amplification and library preparation were performed as described in [47] using the Qiagen Multiplex PCR kit. The 18S amplicon pools were sequenced on an Illumina MiSeq using V3 chemistry and 2 x 300 bp reads. Sequence quality filtering, read merging and primer trimming were performed as described in [34].

A library of 18S sequences for all included arthropod species (from [34]) was used as a reference database to identify the recovered sequences using BLASTn [48], with a minimum e-value of 10^−4^ and a minimum overlap of 95 %. Despite the high raw error rate of nanopore reads, taxonomic status of sequences could be assigned using BLAST, as our pools contained members of highly divergent orders. We compared the qualitative (number of species) and quantitative (abundance of species) recovery of taxa from the communities by nanopore long-read and Illumina short read data. To estimate the recovery of taxon abundances, we calculated a fold change between input DNA amount and recovered reads for each taxon and mock community. A fold change of zero corresponded to a 1:1 association of taxon abundance and read count, while positive or negative values indicated higher or lower read counts than the taxon’s actual abundance.

### MiniBar

We created a de-multiplexing software, called MiniBar. It allows customization of search parameters to account for the high read error rates and has built-in awareness of the dual barcode and primer pairs flanking the sequences. MiniBar takes as input a tab-delimited barcode file and a sequence file in either fasta or fastq format. The barcode file contains, at a minimum, sample name, forward barcode, forward primer, reverse barcode, and reverse primer for each of the samples potentially in the sequence file. The software searches for barcodes and for a primer, each permitting a user defined number of errors, an error being a mismatch or indel. Error count to determine a match can either be a percentage of each of their lengths or can be separately specified for barcode and primer as a maximum edit distance [49]. Output options permit saving each sample in its own file or all samples in a single file, with the sample names in the fasta or fastq headers. The found barcode primer pairs can be trimmed from the sequence or can remain in the sequence distinguished by case or color. MiniBar, written in Python 2.7, can also run in Python 3 and has the single dependency of the Edlib library module for edit distance measured approximate search [50]. MiniBar can be found at https://github.com/calacademy-research/minibar along with test data.

## Results

### Sequencing, specimen recovery and consensus quality

After quality filtering and trimming, our nanopore run yielded 245,433 reads. We tested edit distances of two, three and four bases in MiniBar to demultiplex samples. Increasing edit distances led to a significant increase in read numbers assigned to index combinations (Pairwise Wilcoxon Test, FDR-corrected P-value < 0.05). On average, we found 355 reads per specimen for an edit distance of two, 647 for a distance of three and 1,051 for a distance of four. However, at an edit distance of four, we found a considerable increase of wrongly assigned samples. Using Illumina shotgun sequencing-derived consensus sequences of rDNA from six *Peperomia* plants, we tested the accuracy of the nanopore consensus assemblies based on the three edit distances (Fig. 2). While a distance of four yielded the highest number of assigned reads (1,785 on average), it also led to slightly more inaccurate consensus assemblies, with an average distance of 2.072 % to Illumina based consensus sequences. We found a significant increase of consensus accuracy (Pairwise Wilcoxon Test, FDR corrected *P* < 0.05) for edit distances of two (0.165 % average distance) and three (0.187 % average distance). Despite significant differences in assigned reads (1,091 vs. 637 reads on average), there was not a significant difference in consensus accuracy of edit distances of two versus three bases (Pairwise Wilcoxon Test, FDR corrected *P* > 0.05).

**Figure 2:**
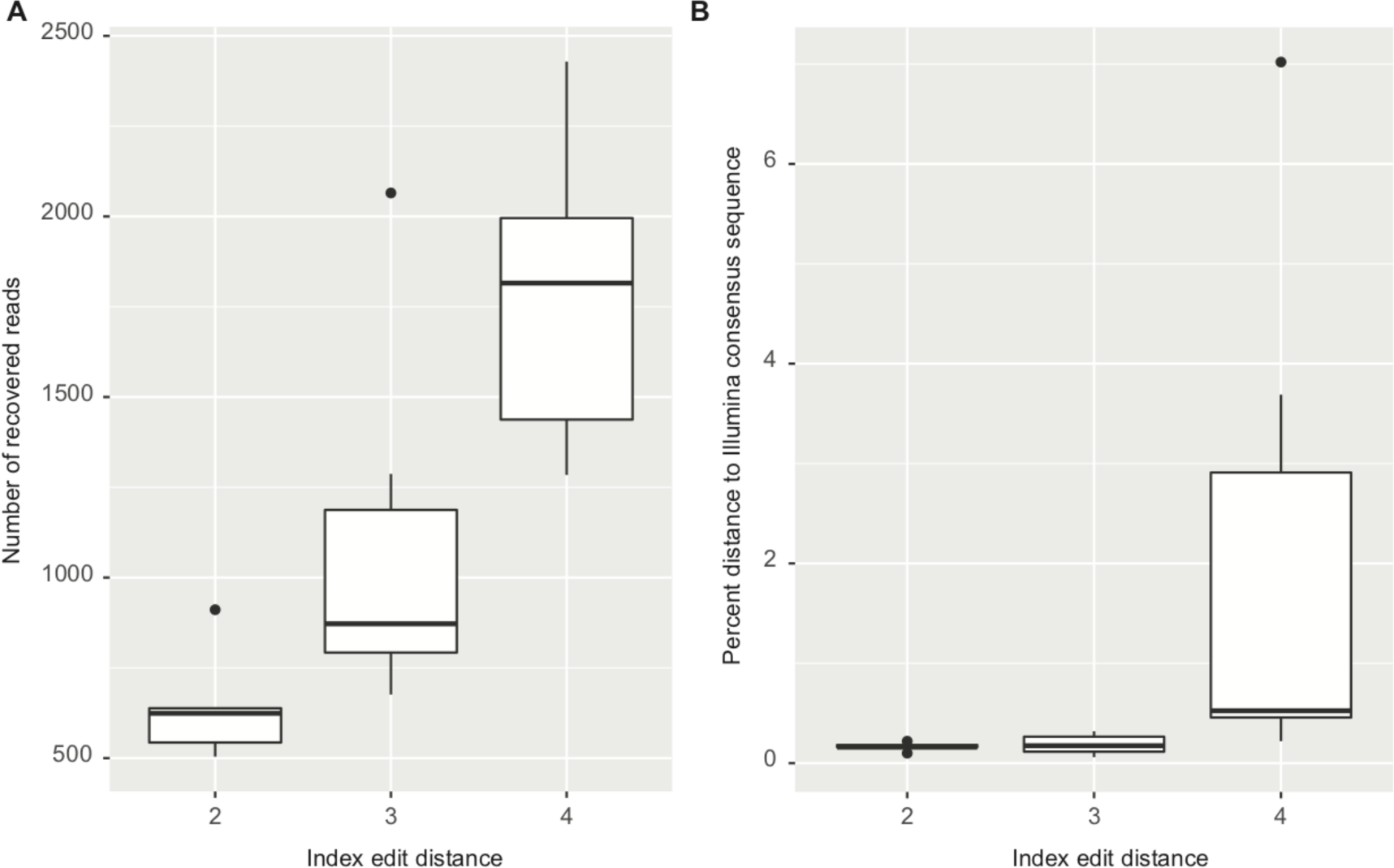
Comparison of recovered sequences and consensus accuracy for different index edit distances in Minibar. A) Number of recovered reads for six *Peperomia* species at index edit distances of two, three and four. B) Pairwise sequence divergence between Illumina and Nanopore based consensus sequences of the same six *Peperomia* specimens at the same index edit distances.

We chose a minimum coverage of 30 (see below) and an edit distance of two (which showed the smallest final consensus error rate) for all subsequent analyses. BLAST analyses suggested a correct taxonomic assignment for the majority of these consensus sequences. However, we found some notable exceptions. For two insect specimens, we amplified mite rDNA sequences. One of these specimens was *Drosophila hydei*, with the mite taxon being a well known phoretic associated with arthropods. A different mite taxon was assembled from an unidentified termite species. A species of isopod and a neuropteran yielded fungal sequences after assembly. The larva of a butterfly and a feeder mealworm *(Zophobas morio)* generated consensus sequences for plants.

A comparison of our consensus sequences for 47 Hawaiian specimens of the spider genus *Tetragnatha* with short Illumina amplicon sequencing-derived 18S and 28S rDNA sequences suggests a very high consensus accuracy. Except for a single specimen, with a single substitution error, all nanopore based consensus sequences were completely identical to the Illumina based consensus. However, the corresponding 18S and 28S fragments did not contain long stretches of homopolymer sequences, where nanopore raw read errors are known to accumulate [51]. Despite containing several homopolymers, the nanopore derived *Peperomia* consensus sequences were highly accurate (Supplementary Fig. 1). Including gaps in the alignment, an average distance of 0.165 % to Illumina based consensus sequences was found. Errors were clustered in Indel regions. After excluding gaps, the average distance dropped to 0.102 %.

We found only a small effect of sequence coverage on consensus assembly accuracy (Supplementary Fig. 2). Even at 10-fold coverage, a low average distance of 0.257% to Illumina consensus sequences was observed. However, at 20-fold coverage, the average distance significantly decreased to 0.128 % (Pairwise Wilcoxon Test, FDR corrected *P* < 0.05). A slight, but not significant, decrease of distance was observed with increasing coverage, with optimal consensus accuracy at 300-fold coverage (0.031 % distance). At coverages larger than 300, the consensus accuracy slightly decreased (average distance of 0.103 % at 800 X coverage).

The length of the rDNA amplicon was quite variable between taxa. Compared to animals, plant specimens showed a significantly shorter amplicon (Pairwise Wilcoxon Test, FDR corrected *P* < 0.05). The length difference was found for the actual gene sequences (18S, 5.8S, 28S: 3,063 vs 2,781 bp on average; Supplementary Fig. 3A) as well as including the ITS sequences (3,741 vs. 3,241 bp on average, Supplementary Fig. 3B). Within arthropods, we found significant length differences between arachnid and insect sequences. On average, insects carried a significantly longer rDNA sequence than arachnids (Supplementary Fig. 4; Pairwise Wilcoxon Test, FDR corrected *P* < 0.05). This holds true for the gene sequences (3,154 vs. 3,047 bp for 18S, 5.8S, 28S on average), as well as the whole amplicon, including ITS sequences (4,192 vs. 3,644 bp on average). While most spiders showed very stable length distributions for the rDNA amplicon length (average length ± standard deviation across all Araneae: 3,629 bp ± 81), several insect orders had rDNA sequences of more variable length (Coleoptera: 4,488 bp ± 352; Lepidoptera: 4363 bp ± 603).

**Figure 3.**
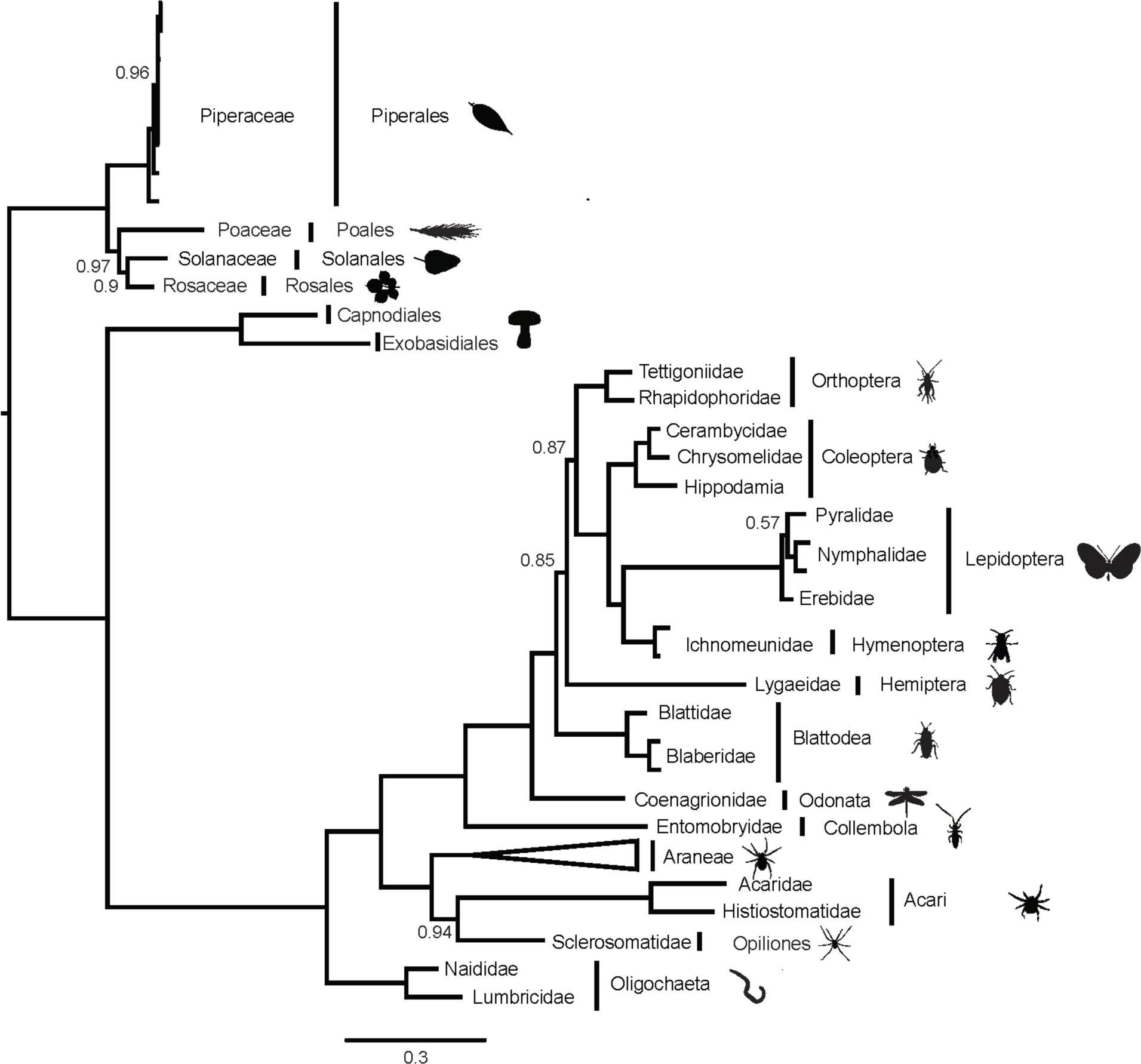
Bayesian consensus phylogeny based on a 3,656 bp alignment of 18S, 5.8S and 28S sequences of 117 animal, fungal and plant taxa. The phylogeny is rooted using plants as outgroup. Branches are annotated with family and order level taxonomy. The Araneae clade of 83 specimens is collapsed. Only posterior probability values below 1 are displayed.

In contrast to the variable length of the rDNA cluster, we found a very stable GC content across the whole taxonomic spectrum (46.75 ± 2.67 % across all taxa). GC content of plants and animals was highly similar (Supplementary Fig 3c) (plants: 46.01 ± 1.66 %; animals: 46.82 ± 2.74 %). Highly similar GC content was also found between insects (46.67 ± 3.73 %) and arachnids (46.93 ± 2.47 %) (Supplementary Fig 4c).

### Phylogenetic reconstruction

We generated an alignment of 3,656 bp for 117 concatenated 18S, 5.8S and 28S sequences of plants, fungi, annelids and arthropods. Our phylogeny was well supported (most posterior support values equal one; Fig. 3). A basal split separated plants from fungi and animals. Within plants, the genus *Peperomia* was recovered as monophyletic. Fungi formed the sister group of animals. Within animals, annelids formed a separate clade from arthropods. Arthropods separated into arachnids and hexapods. Each separate arthropod order formed well supported groups. The hexapod phylogeny generally resembled that found in latest phylogenomic work [52]. The Collembola species *Salina* sp. formed the base to the insect tree, followed by the odonate *Argia* sp. A higher branch led to Blattodea, Hemiptera and Orthoptera. However, the support values for the relationships between these three orders were comparatively low (~ 0.85). Finally, holometabolan insects (Hymenoptera, Coleoptera and Lepidoptera) were recovered as monophyletic. The two Acari species, together with Opiliones, formed the sister clade to the monophyletic Araneae clade.

Next, we generated a separate alignment of rDNA sequences for 83 spiders, including both ITS regions (totaling 4,214 bp). The spider phylogeny was also strongly supported (Fig. 4). Overall, our phylogenetic tree topology agreed with the most recent phylogenetic work of [53] and [35]. With the haplogyne *Segestria* sp. (family Segestriidae) forming the root, we recovered the so-called RTA clade (represented in our dataset by families Agelenidae, Amaurobiidae, Anyphaenidae, Cybaeidae, Desidae, Eutichuridae, Lycosidae, Philodromidae, Psechridae, Salticidae and Thomisidae) and the Araneoidea (Araneidae, Linyphiidae, Tetragnathidae, Theridiidae) as two well supported monophyla. Within these clades, all families and genera formed well supported monophyletic groups. Similar to recent studies, we found the Marronoid clade as basal to the rest of the RTA clade; more derived clades were the Oval Calamistrum and the Dionycha clade. Inter-family relationships also closely matched those found in recent work: Lycosidae was basal to the clade formed by Psechridae and Thomisidae; Salticidae was closest to Eutichuridae and Philodromidae, with Anyphaenidae falling basal within Dionycha. Within Araneoidea, our results differed slightly from recent studies in that we recovered Tetragnathidae, rather than Theridiidae, as basal.

**Figure 4.**
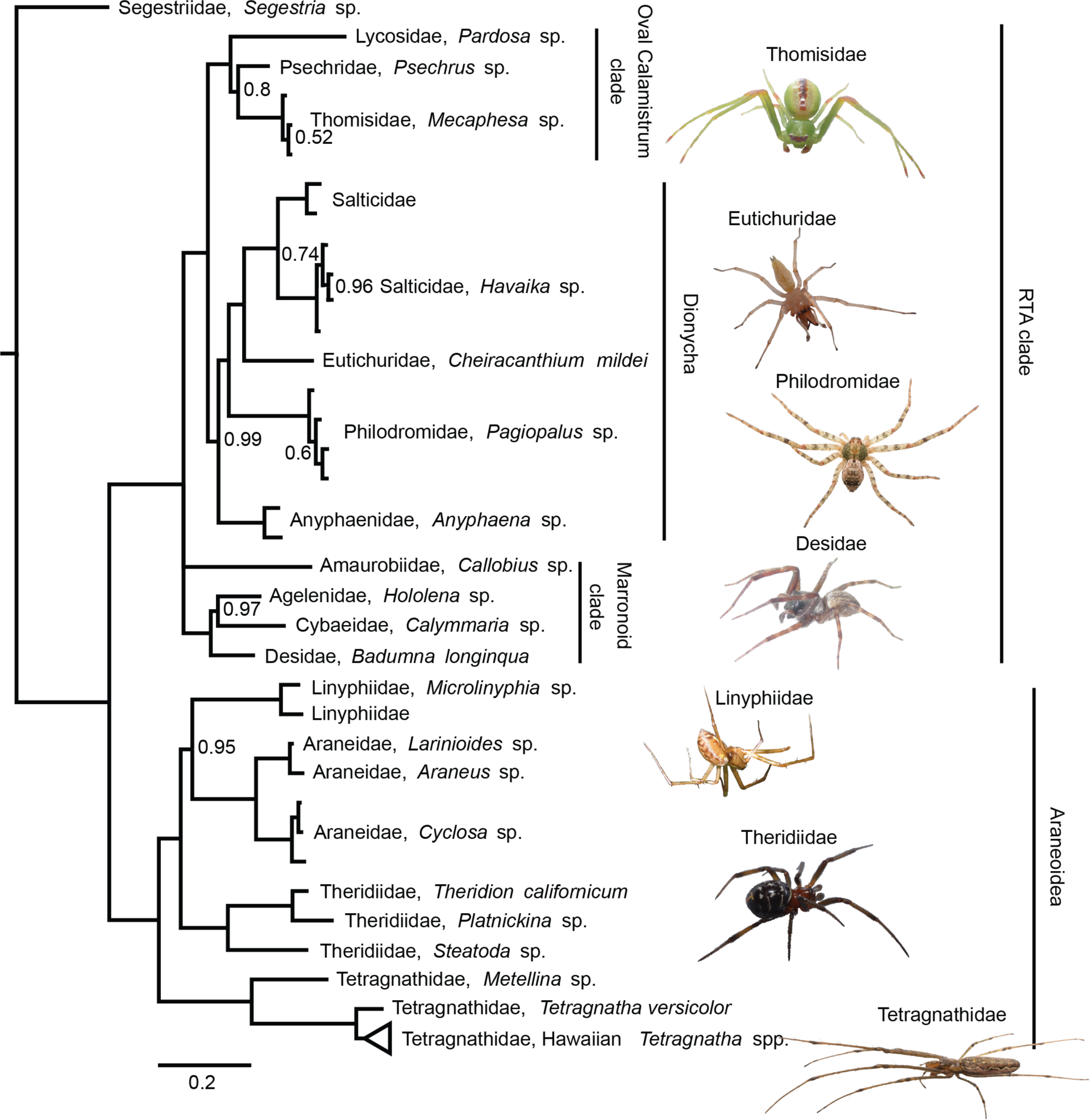
Bayesian consensus phylogeny of 83 spiders in 16 families, based on a 4,214 bp alignment of 18S, ITS1, 5.8S, ITS2 and 28S. The phylogeny is rooted using the basal haplogyne *Segestria* sp. The clade containing Hawaiian members of the genus *Tetragnatha* is collapsed (the uncolapsed clade is shown in Fig. 5). Only posterior probability values below 1 are displayed.

We recovered Hawaiian *Tetragnatha* as a well supported monophyletic clade within the Tetragnathidae. We found two main clades of Hawaiian *Tetragnatha* (Fig. 5), both of which have been supported by earlier work [54–57]: the orb weaving clade and the “Spiny Leg clade” of actively hunting species. All *Tetragnatha* species formed monophyletic groups, and the relationships among different species were mostly well supported. Within the Spiny Leg clade, species fell into one of four ecotypes, each of which is associated with a particular substrate type: “large brown” *(T. quasimodo)* with tree bark, “small brown” *(T. anuenue, T. obscura* and *T. restricta)* with twigs, “green” *(T. brevignatha* and *T. waikamoi)* with green leaves, and “maroon” *(T. perreirai* and *T. kamakou)* with lichen. While green and maroon ecotypes clustered phylogenetically, small brown species appeared in three separate clades on the tree. Within the orb weaving clade, *T. hawaiensis*, a generalist species which occurs on all of the Hawaiian Islands, fell basal. The characteristic web structures of some of these species have been documented [35, 58]. We found a pattern of apparent convergence in web structure for some species. *T.* sp. “emerald ovoid” spins a loose web with widely spaced rows of capture silk. *T. hawaiensis* and *T.* sp. “eurylike,” which are distant relatives within the Hawaiian *Tetragnatha* clade, both spin webs of medium silk density, i.e. with more rows of capture silk per unit area than *T.* sp. “emerald ovoid.” *T. perkinsi* and *T. acuta* each spin a web structure that is not comparable in its silk density or size to any other known *Tetragnatha* species in this group [58], and are thus classified as “unique”.

**Figure 5.**
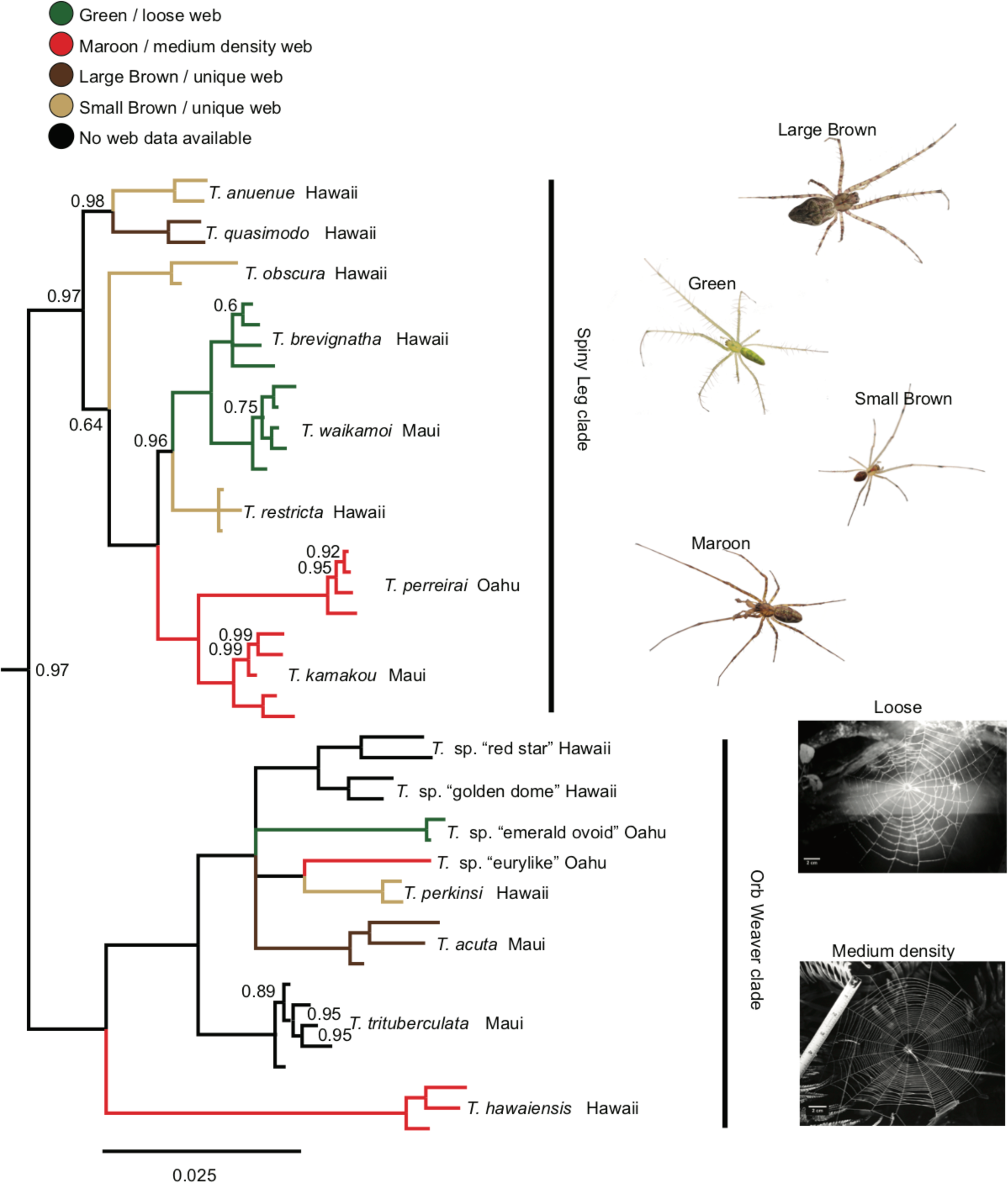
Section of the same phylogeny as Fig. 4, with expansion of the clade of 16 Hawaiian *Tetragnatha* species. Different “Spiny Leg” ecomorphs and web architectures are indicated by branch coloration. Only posterior probability values below 1 are displayed.

Our inferred genetic distances for rDNA sequences within and between Hawaiian *Tetragnatha* species were significantly correlated to those found for COI sequences of the same taxa (R^2^ = 0.70, *P* < 0.001) (Fig. 6a). A Mantel test also suggested highly significant correlation of mitochondrial COI and nuclear rDNA based distances (Mantel test, 9999 replicates, *P* < 0.001). Hence, the rDNA cluster supported a very similar pattern of genetic differentiation to COI. However, the faster evolutionary rate of COI was reflected in lower distances for the whole rDNA than for COI. Interspecific distances were significantly higher than intraspecific ones for COI and rDNA (Fig 6b,c). No overlap of intra and interspecific distances was evident for COI, suggesting the presence of a barcode gap. A small overlap of intra and interspecific distances was evident for the rDNA (Supplementary Table 3). Like the combined rDNA cluster, genetic distances for different parts of the rDNA cluster all showed significant correlation with COI based distances, when analyzed separately (R^2^ 28S = 0.57, R^2^ ITS1 = 0.68, R^2^ ITS2 = 0.56, *P* < 0.001) (Supplementary Fig. 5). While the 28SrDNA showed considerably lower distances than COI, those for ITS1 and ITS2 were more comparable to COI (Supplementary Fig. 5b-d). Yet, interspecific and intraspecific distances for COI were significantly different from those for any part of the rDNA cluster (Pairwise Wilcoxon Test, FDR corrected *P* < 0.05).

**Figure 6.**
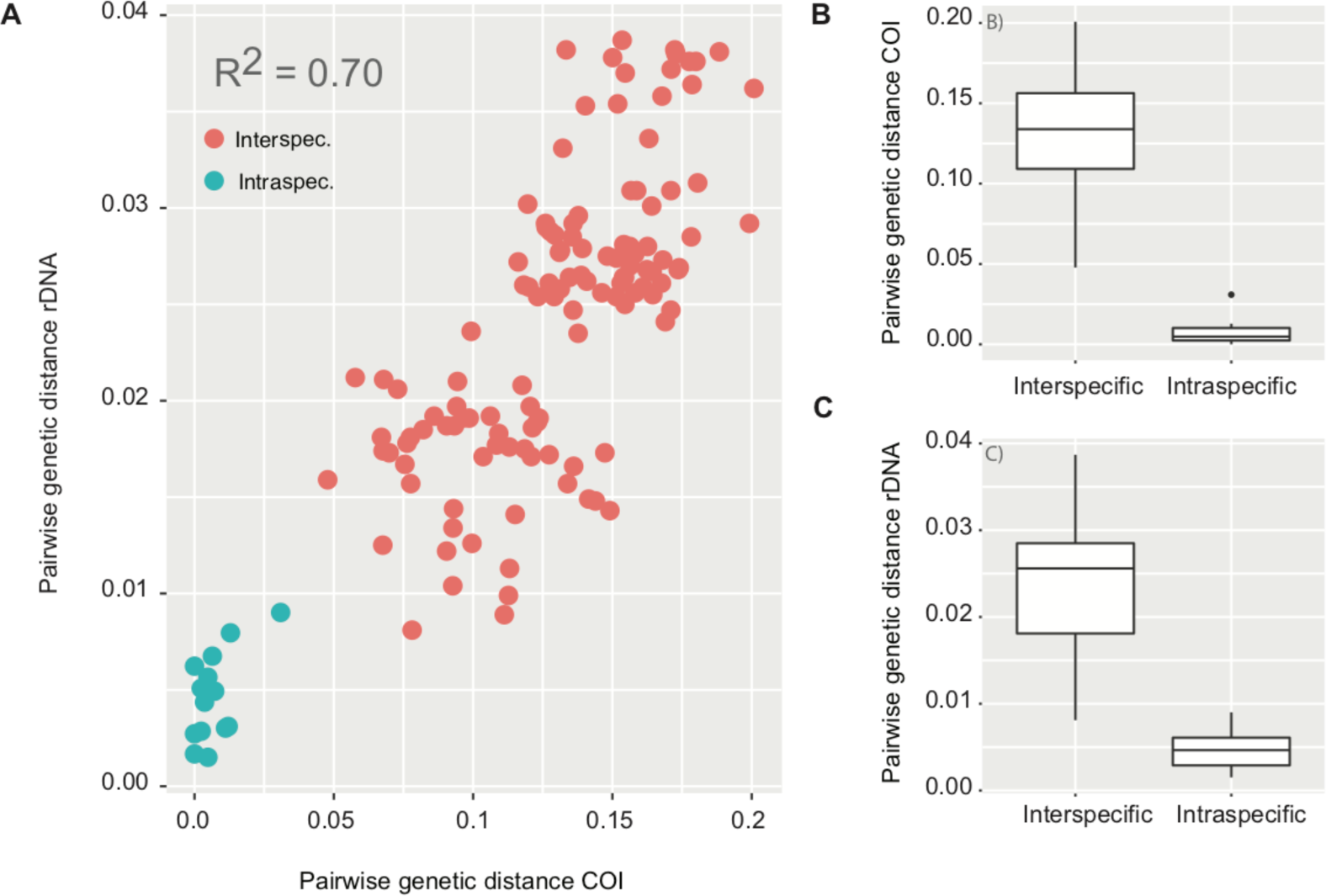
Inter and intraspecific genetic distances for the nuclear rDNA and mitochondrial COI for Hawaiian *Tetragnatha* **spiders.** A) Correlation of pairwise genetic distance between (red) and within (green) 16 Hawaiian *Tetragnatha* species based on COI and the full rDNA amplicon. B) Interspecific and intraspecific genetic distances for the same spider species based on mitochondrial COI and C) the whole rDNA amplicon.

### Field trial in the Amazon rainforest

On March 26, 2018 we set out to test this method and a portable laboratory (as described in Pomerantz, et al. [25]) during an expedition to the Peruvian Amazon at the Refugio Amazonas Lodge (Supplementary Fig. 6). This field site is a “Terra firme” forest in the sector of “Condenado”, approximately two and a half hours by boat up river from the native community of Infierno on the buffer zone of the Tambopata National Reserve. We collected plant and insect material, extracted DNA, amplified the rDNA cluster, and sequenced material on the MinION platform using the MinKNOW offline software (provided by ONT). The first run generated 17,149 reads and the second one 20,167 reads. We generated consensus sequences for five out of the seven analyzed specimens. One plant sample and the grasshopper could not be assembled due to too low read coverage. Moreover, BLAST analysis of the reads assigned to the grasshopper suggested that we had sequenced a mite, instead of the grasshopper DNA. The unidentified insect eggs resulted in a butterfly consensus sequence, possibly a Pierid species.

### Nanopore based arthropod metabarcoding

On average, we recovered 2,645 reads for each Illumina sequenced mock community and 1,149 for each nanopore mock community. The optimized Illumina amplicon sequencing based 18SrDNA protocol resulted in a very good taxon recovery. All nine taxa were recovered from all four mock communities (Fig. 7). Moreover, the Illumina based protocol allowed for accurate predictions of taxon abundances. The average fold change between input DNA and recovered read count was closely distributed around zero (Supplementary Fig 6). In contrast, the long-read nanopore protocol showed very biased qualitative and quantitative taxon recovery (Fig. 7). On average, only 83.33 % of taxa were recovered per nanopore sequenced mock community. Moreover, the fold change of input DNA and recovered read count were highly biased between taxa. Some taxa were considerably over or underrepresented among the read population. This led to a significantly higher variation of fold change between input DNA and read count compared to the Illumina amplicon based protocol (Levene’s test *P* < 0.05; Supplementary Fig. 7). A reduction of PCR annealing temperature did result in a considerable increase of Odonata sequences, but overall did not have a strong effect on qualitative (77.78 % of taxa recovered) or quantitative taxon recovery (Fig. 7). The variation of fold change between different PCR annealing temperatures was not significantly different (Levene’s test, *P* > 0.05). A reduction of PCR cycle number by 10 also did not yield any significant effect on qualitative (88.89 % of taxa recovered) or quantitative taxon recovery (Supplementary Fig. 7).

**Figure 7:**
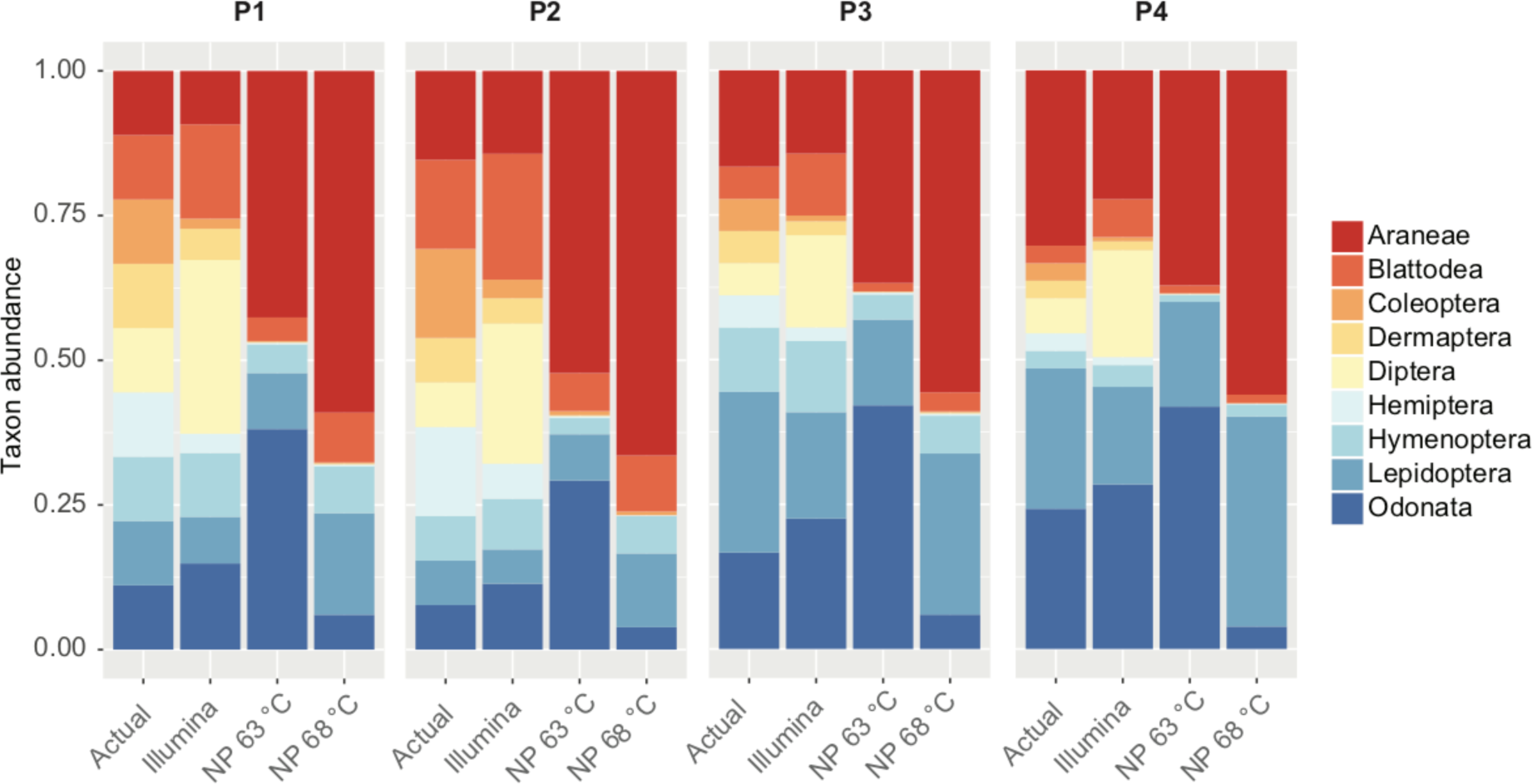
Relative abundances for nine arthropod species in our four mock communities (actual), compared to an Illumina amplicon sequencing protocol, and nanopore protocols at 63 °C and 68 °C annealing temperature

## Discussion

### Phylogenetic and taxonomic utility of long rDNA amplicons

Developments in long-read sequencing hold great promise for molecular taxonomy and phylogenetics across very broad taxonomic scales. We recovered phylogenetic relationships across the eukaryote tree of life, which were mostly consistent with the current state of research (e.g. [52]). Separate orders of arthropods all formed well supported monophyletic groups. Our spider phylogeny was highly congruent with recent work based on whole transcriptomes [35] and multi-amplicon data [53]. Moreover, using the rDNA cluster allowed us to resolve young phylogenetic divergences: the relationships within the recent adaptive radiation of the genus *Tetragnatha* in Hawaii confirmed previous research [59, 60].

Besides their high phylogenetic utility, long rDNA amplicons showed excellent support for taxonomic hypotheses. All morphologically identified species of Hawaiian *Tetragnatha* were recovered as monophyletic groups. The divergence patterns and taxonomic classifications of spiders based on rDNA were strongly correlated to those based on mitochondrial COI, the most commonly used animal barcode marker [4]. rDNA may thus be ideal to complement mitochondrial barcoding. A universal and variable nuclear marker as a supplement to COI barcoding will be particularly useful in cases of mito-nuclear discordance due to male biased gene flow [10, 61], hybridization [12] or infections with reproductive parasites [11].

Their high phylogenetic utility across very broad taxonomic categories also provide long rDNA amplicons with a distinct advantage over short read barcoding protocols, which are not well suited to support broad scale phylogenetic hypotheses [62]. The inclusion of long amplicons would make it possible to scale up barcoding from simple taxon assignment to community wide phylogenetic inferences [9]. Recently, the amplification of whole mitochondrial genomes was suggested for animal barcoding [63]. This would increase taxonomic and phylogenetic resolution and thus alleviate some disadvantages of short COI amplicons. However, it is challenging to develop truly universal primers to target mitochondrial genomes across a wide range of taxonomic groups [64]. Moreover, mitochondrial genomes will not allow cases of mito-nuclear discordance to be identified. A straightforward way to achieve highly resolved phylogenies may be the combination of long rDNA amplicon sequencing with multiplex PCRs of short mitochondrial amplicons, to amplify multiple mitochondrial DNA fragments [65].

### Simple, accurate, universal and cost efficient long-read DNA barcoding

Despite the high raw read error of nanopore data, consensus sequences were highly accurate, and library preparation and sequencing for our protocol are simple and cost efficient. Using a single pair of universal primers, long rDNA amplicons can be amplified across diverse eukaryote taxa. A simple dual indexing approach during PCR allows large numbers of samples to be pooled before library preparation [27]. Only a single PCR is required per specimen, while subsequent cleanup and library preparation can be performed on pooled samples. The simplicity of our approach is additionally highlighted by its effectiveness even under field conditions in a remote rainforest site. Nanopore sequencing technology is affordable and universally available to any laboratory. Our ONT MinION generated about 250,000 reads per run. Aiming for about 1,000 reads per amplified specimen, 250 long rDNA barcodes could be generated in single MinION run. Input DNA amounts for different specimens will have to be carefully balanced to maximize the recovery. The total reagent costs, including PCR, library preparation and sequencing, then amount to less than $4 for each long barcode sequence generated.

### Pitfalls of nanopore based long-read barcoding

While our protocol was generally straightforward and reliable, we found several drawbacks, which require further considerations and optimization. First, it needs to be noted that long rDNA amplification will not be possible with highly degraded DNA molecules, e.g. from historical specimens [66]. Moreover, amplification success of long range PCRs proved less consistent than that for amplification of short amplicons. We observed a complete failure of some PCRs when too high template DNA concentrations were loaded. The long range polymerase may be more sensitive to PCR inhibitors present in some arthropod DNA extractions [67]. PCR conditions will have to be carefully optimized for reliable and consistent amplification. We also found that highly universal eukaryote primers may result in undesired amplification, for example plants from beetle and butterfly larval guts, phoretic mites, or fungal sequences. However, as long as the DNA of the target taxon is still dominating the resulting amplicon mixture, this undesired amplification will not affect consensus calling. It may be advisable to check the taxonomic composition of amplicon samples before assembly, e.g. by blasting against a reference library. To avoid unspecific amplification, PCR primers could also be redesigned to exclude certain lineages from amplification. It should also be noted that our approach results in only a single consensus sequence for each processed specimen. As a diploid marker, the rDNA cluster can contain heterozygous positions in some specimens, in particular within the ITS regions. This information is currently lost, and a different assembly approach may be necessary to recover heterozygosity as well. Furthermore, index length and edit distance are also important considerations. We used indexes of 15 bp and with a minimum distance of 10 bp to index both sides of our amplicons. Index edit distance of only 4 bp between samples already led to considerable cross-specimen index bleeding. It may thus be better to increase the length and edit distances of indexes. Indexes of 20 or 30 bp could be easily attached to the 5’-tails of PCR primers without strongly affecting PCR efficiency.

### Nanopore based arthropod metabarcoding

It is well known that Illumina amplicon sequencing of short 18SrDNA fragments can yield very accurate qualitative and quantitative taxon recovery in metabarcoding experiments [48], a finding that is confirmed by our results. In contrast, little is known on the performance of long-read nanopore sequencing for community diversity assessments [32]. Our long barcode based approach resulted in the dropout of several taxa and highly skewed relative taxon abundances. Skewed abundances were already found in microbial community analysis using nanopore [32]. In the simplest case, primer mismatches may be responsible for biased amplification [32, 68]. However, the targeted priming sites in our study were extremely conserved. Also, a change of PCR cycle number and annealing temperature did not have a strong effect on taxon abundances, as would be expected in the case of PCR priming bias [69]. Another possibility is the preferential amplification of template molecules with a certain GC content by the DNA polymerase [33]. However, we found the GC content of the rDNA cluster to be very stable across taxa. Yet another potential explanation for the differential recovery of taxa in community samples is taxonomic bias in DNA degradation [70], but we do not expect DNA degradation to have played a role in our experiment because we used only high quality DNA extractions (verified by gel electrophoresis) from fresh specimens. The most plausible explanation appears to be that variable rDNA lengths are found between different taxa. It is well known that shorter sequences are amplified preferentially in a PCR, especially after it reaches the plateau stage [71]. Such dominance of shorter amplicons could explain the observed biases very well. In fact, the most abundant taxon in our pools was a spider, which also had the shortest amplicon length. The dominant amplification of shorter sequences may also explain the amplification of plant DNA from a butterfly and a flour beetle larva, as plants showed considerably shorter rDNA amplicons than insects. We found a very high variation of rDNA amplicon length within many taxonomic groups, this could be a considerable problem for long read metabarcoding applications. More research into the causes and possible mitigation of these biases will be required before long-read sequencing can be routinely utilized for metabarcoding applications.

## Conclusion

Sequencing long dual indexed rDNA amplicons on Oxford Nanopore Technologies’ MinION is a simple, cost effective, accurate and universal approach for eukaryote DNA barcoding. Long rDNA amplicons offer high phylogenetic and taxonomic resolution across broad taxonomic scales from kingdom down to species. They also prove to be an excellent complement to mitochondrial COI based barcoding in arthropods. However, despite the long-read advantages in the analysis of separate specimens, we found considerable biases associated with sequencing bulk community samples. The observed taxonomic bias is possibly a result of taxon-specific length variation of the rDNA cluster and preferential amplification of species with shorter rDNA. Further research into the sources of the observed bias is required before long rDNA amplicon sequencing can be utilized as a reliable resource for the analysis of bulk samples.

## Author contributions

HK and SP designed the study. HK, AP, SRK, JYL, VS and JDS collected the specimens. Laboratory work was carried out by HK, AP and SRK and the data were subsequently analyzed by HK, AP, JBH, SRK and SP. The paper was writen by HK, AP, JBH, SRK, JYL, VS, JDS, NHP, RGG, SP.

## Acknowledgements

We thank Taylor Liu for help during laboratory work, and Natalie Graham and Tara Gallant for help during specimen collection. Hitomi Asahara graciously provided access to a laboratory facility and the necessary software for our MinION sequencing run. We thank the State of Hawaii Department of Land and Natural Resources and the Servicio Nacional Forestal y de Fauna Silvestre, who provided collection permits, Anna Holmquist for providing the *Psechrus* sp. specimen, rainforest Expeditions and Gabriela Orihuela for providing assistance and support with fieldwork in Peru.

